# Automatic Geometry-based Estimation of the Locus Coeruleus Region on T_1_-Weighted Magnetic Resonance Images

**DOI:** 10.1101/2024.01.23.576958

**Authors:** Iman Aganj, Jocelyn Mora, Bruce Fischl, Jean C. Augustinack

## Abstract

The locus coeruleus (LC) is a key brain structure implicated in cognitive function and neurodegenerative disease. Automatic segmentation of the LC is a crucial step in quantitative non-invasive analysis of the LC in large MRI cohorts. Most publicly available imaging databases for training automatic LC segmentation models take advantage of specialized contrast-enhancing (e.g., neuromelanin-sensitive) MRI. Segmentation models developed with such image contrasts, however, are not readily applicable to existing datasets with conventional MRI sequences. In this work, we evaluate the feasibility of using non-contrast neuroanatomical information to geometrically approximate the LC region from standard 3-Tesla T_1_-weighted images of 20 subjects from the Human Connectome Project (HCP). We employ this dataset to train and internally/externally evaluate two automatic localization methods, the Expected Label Value and the U-Net. We also test the hypothesis that using the *phase* image as input can improve the robustness of out-of-sample segmentation. We then apply our trained models to a larger subset of HCP, while exploratorily correlating LC imaging variables and structural connectivity with demographic and clinical data. This report contributes and provides an evaluation of two computational methods estimating neural structure.

## 1 Introduction

The locus coeruleus (LC) is a small elongated hyperpigmented nucleus in the rostral pontine brainstem (1) It synthesizes most of the brain’s norepinephrine (2) and is involved in various cognitive functions (3). The LC undergoes neuron loss in the early stages of many neurodegenerative diseases (4-6), such as Alzheimer’s disease (7-9) and Parkinson’s disease (10, 11) through the accumulation of tau pathology (12, 13) and α-synuclein (14), respectively. Non-invasive assessment of the LC integrity *in vivo*, namely via magnetic resonance imaging (MRI), helps to elucidate how LC degeneration relates to the progression and symptoms of neurodegenerative diseases (4-6). Patterns of structural connectivity of the LC to other brain regions – quantified via diffusion MRI (dMRI) (15, 16) – may further inform us about the pathology distribution in the brain, particularly in the context of Alzheimer’s disease, where some hypothesize that tau protein may transmit neuron to neuron from the LC to other areas (17).

Quantitative analysis of the LC from MRI requires knowledge of the LC location. Manual annotation of the LC in a large dataset not only necessitates significant expert effort, but yields a precision limited by moderate inter- and intra-rater variability (18). Automatic LC localization (19-22), which is not yet widely available in conventional neuroimaging toolboxes, is therefore highly desirable, as it can facilitate large-scale imaging studies that would have the power to detect subtle changes in the LC in health and disease.

To enhance the contrast of the LC in the MR image, the high concentration of neuromelanin in the LC (23) has been exploited. To that end, several neuromelanin-sensitive MRI sequences (24) have been successfully employed, including the T_1_-weighted (T_1_W) Turbo Spin Echo (TSE) (18, 20, 25-27) and the magnetization transfer (13, 22, 28-31) sequences. The enhanced LC contrast on images acquired with such sequences allows for manual delineation of the LC and the creation of datasets that include gold-standard LC labels. Using a dataset like this for training, a supervised automatic segmentation algorithm can segment the LC on a new similar-contrast image.

Neuromelanin-sensitive MRI, however, typically has a high specific absorption rate (5, 32, 33), and may also be suboptimal for younger adults due to their lower neuromelanin levels (6, 30, 34). Consequently, such a sequence is often not included in large open-access MRI databases of healthy or diseased populations. As for standard MR images (e.g., T_1_W images included in almost all MRI databases), the boundaries of the LC cannot be delineated on these images due to the lack of contrast; therefore, the location of the LC can only be approximated at best relative to its surrounding structures using prior neuroanatomical information. Such geometrical localization of the LC might still be useful for some subsequent analyses not requiring high label accuracy. For instance, deep neural networks trained on silver-standard labels have shown to be capable of producing more reliable segmentation results than the labels they were trained on (35).

Several LC atlases are publicly available (16, 18, 26, 27, 30, 31, 36, 37), which can be employed to automatize the localization of the LC via atlas alignment and label propagation. The original datasets used to create these public atlases, however, are not generally available. As a result, users cannot apply other supervised segmentation methods to localize the LC, such as those based on modern deep neural networks (38). A public database of approximate LC region masks accompanied with corresponding MR images with standard (e.g., T_1_W, rather than neuromelanin-sensitive) contrasts is thus desirable. Such a database could help the research community to use existing or new methods to develop automatic LC localization tools that are applicable to many available databases, thereby facilitating large-scale retrospective and prospective analyses involving the LC.

Our contributions in this work are as follows:

- We first manually approximate the LC region on the 3T T_1_W images of 20 subjects of the open-access *Human Connectome Project (HCP)* (39), sharing the geometrically annotated masks with the research community (see Section 8). To our knowledge, other publicly available datasets with manual LC labels instead contain 3T-TSE/7T T_1_W images (33) or functional MR images (40). Our geometrical estimation is based on dimensional (instead of contrast) information and emphasizes the sensitivity of the detection, thereby resulting in LC masks slightly larger than the LC (i.e., encompassing the LC and some surrounding area).
- We then train two automatic segmentation methods of Expected Label Value (ELV) (41) and U-Net (38) on the abovementioned dataset, evaluating the LC localization ability internally as well as on an external dataset (33).
- Inspired by a previous observation (41), we test the hypothesis that using the phase image (i.e., discarding the magnitude Fourier data) would improve the performance of the above models on external datasets (data from different sources).
- We finally apply a trained model to one hundred HCP subjects and analyze the volume, image intensity, and dMRI structural connectivity of the LC masks, correlating them with non-MRI variables.

In the following, we will describe our methods (Section 2), provide our results (Section 3), and discuss them (Section 4).

## 2 Method

### 2.1 Manual LC Region Estimation

The human LC is a thin and long column of neurons that extends through multiple levels of the brainstem (1, 5). Located in the rostral pons, the LC is on average 14.5 mm long and 2-2.5 mm wide (1). Due to the lack of LC contrast on T_1_W images, we instead used these dimensional landmarks collectively to approximate the LC location: 3 mm lateral from the midline, 1 mm rostral to the fourth ventricle, and 16-20 mm above the pontomedullary junction.

We used Freeview of FreeSurfer (42) to manually create masks of the areas containing each of the left and right LCs on preprocessed 3T T_1_W MPRAGE images (`T1w_acpc_dc_restore_brain.nii.gz`) of the first 20 subjects of the “100 Unrelated Subjects” group of the HCP (39), which had the isotropic voxel size of (0.7 mm)^3^. Our localization approach prioritizes sensitivity to specificity (i.e., includes more voxels than the LC alone), producing masks that are somewhat inflated compared to the actual LC boundaries.

### 2.2 Automatic LC Region Estimation

#### 2.2.1 Approaches

We trained two supervised image segmentation methods, both implemented in MATLAB, on our 20-subject dataset to automatically approximate the presumptive left or right LC areas (separately), as described below. We binarized the output soft mask and retained the largest connected component.

We used the ELV supervised segmentation (41) (see Section 8 for toolbox) as our first method, which creates a fuzzy map from a combination of labels suggested by all atlas-to-image transformations, weighted by a measure of transformation validity (without explicit deformable registration). The ELV method inherently uses *phase* images as input (obtained by computing the Fourier transform of the image, discarding the magnitude data, and computing the inverse Fourier transform), which we call “ELV [phase]”. The map can also be modulated by an image intensity prior (41), i.e. “ELV [phase + image]”, to benefit from the image intensity information initially excluded from the phase data.

The second method we used was the U-Net neural network architecture (38), which is a convolutional neural network consisting of a contracting path to capture context, a symmetric expanding path for precise localization, and cross connections. We employed a U-Net with 2 down-sampling layers and 16 initial filters (at the first convolutional layer), with the Dice coefficient as the objective function. We used the Adam optimizer to train the network for 20 epochs on 3D sample patches of size 132×132×132 with a mini-batch size of 8. We initialized the learning rate as 0.002 and dropped it by 95% every 5 epochs. The test subject’s LC region was then predicted by averaging the label scores of overlapping patches (stride 10).

#### 2.2.2 Validation

For performance evaluation, we first internally assessed the automatic localization via leave-one-out cross-validation (i.e., trained on 19 images and tested on the remaining image, repeating it for 20 test images). We compared the automatically generated mask with the manual one using the Dice similarity coefficient as the evaluation metric.

For external (out-of-sample) validation, we then applied our models (that had been pretrained on the 20 HCP subjects) to 12 7T T_1_W images from the previously unseen dataset by Tona *et al* (33), which had LC labels manually delineated from 3T T_1_W TSE images. The images from the latter dataset had the voxel size of 0.70 × 0.64 × 0.64 mm^3^, thereby requiring resampling to match the HCP resolution of (0.7 mm)^3^. Since we trained our models on brain-masked HCP data, we extracted the brain in the new dataset using the SPM12 software package (43).

Note that our geometrical LC approximation produces a generous area containing the LC, and, as such, comparing it to the specific label of the actual LC in external validation results in a suboptimal Dice score. For instance, if our localized area has a volume α times larger than that of the LC label, then the Dice will be no higher than 2⁄(1 + α). Nonetheless, this comparison can still help to assess how much our automatic inflated LC neighborhood overlaps with the LC.

#### 2.2.3 Phase Image as Input

We have previously observed that ELV [phase + image] outperformed ELV [phase] in internal cross-validation, but not in external out-of-sample validation (41). We hypothesized that, being less sensitive to inter-dataset variation in image intensity, the phase image might be more robust to domain shift, resulting in better model performance than the image itself would in external validation. We tested this by comparing the two abovementioned ELV variations, as well as comparing the original U-Net (“U-Net [image]”) to a variation of it that received the phase image as input (“U-Net [phase]”) and a two-channel-input variation that received both the image and the phase as input (“U-Net [image + phase]”).

Prompted by the different image intensity distributions of our two datasets, we also experimented with normalizing the input image by its intensity standard deviation during both training and testing (denoting the normalized image with “nmz”), which led to additional variations such as “ELV [phase + nmz]”, “U-Net [nmz]”, and “U-Net [nmz + phase]”. For the latter two U-Net variations, the input layer also performs patch-wise zero-meaning and normalization.

### 2.3 Analysis of HCP Data

We applied the default implementation of the U-Net to the “100 Unrelated Subjects” group of the HCP. We used the resulting left and right LC masks to compute their volumes as well as the mean T_1_W and T_2_W image intensities inside them. We then propagated the masks to the dMRI space, and similarly computed the mean fractional anisotropy (FA) and the mean apparent diffusion coefficient (ADC), resulting in a total of 10 LC (regional) imaging variables.

Next, we performed an exploratory analysis to correlate the imaging variables with 504 non-MRI variables (demographics, medical history, family history, dementia/cognitive exam scores, personality/emotion tests, motor/sensory tests, task performance, etc.). We Bonferroni-corrected the Pearson’s correlation *p*-values for multiple comparisons through multiplication by the numbers of imaging and non-MRI variables, i.e., *p*_*B*_ = *p* × 10 × 504. We visually inspected relationships with *p*_*B*_ < 0.05 to exclude any spurious correlations due to outliers (e.g., avoiding situations with most data points clustered together with no obvious relationship), reporting the surviving significant correlations.

Finally, we examined the associations between the connectivity pattern of LC and non-MRI variables by quantifying dMRI-derived structural connectivity of the LC area to the rest of the brain. We used our open-source toolbox (see Section 8) to reconstruct the diffusion orientation distribution function in constant solid angle (44), perform Hough-transform global probabilistic tractography (45), compute a symmetric structural connectivity matrix between the two LC areas and 85 other brain regions segmented by FreeSurfer (42), and augment the raw matrices with indirect connections (46). We have previously described this pipeline in detail (47).

## 3 Results

### 3.1 Manual LC Region Estimation

We have provided our geometrically annotated (enlarged) LC masks for the 20 HCP subjects to the public (Section 8). As expected, we did not observe any LC contrast on T_1_W images to guide the manual delineation of the LC region. Figure 1 shows the approximated LC masks (in green/yellow) for a representative subject.

**Figure 1.**
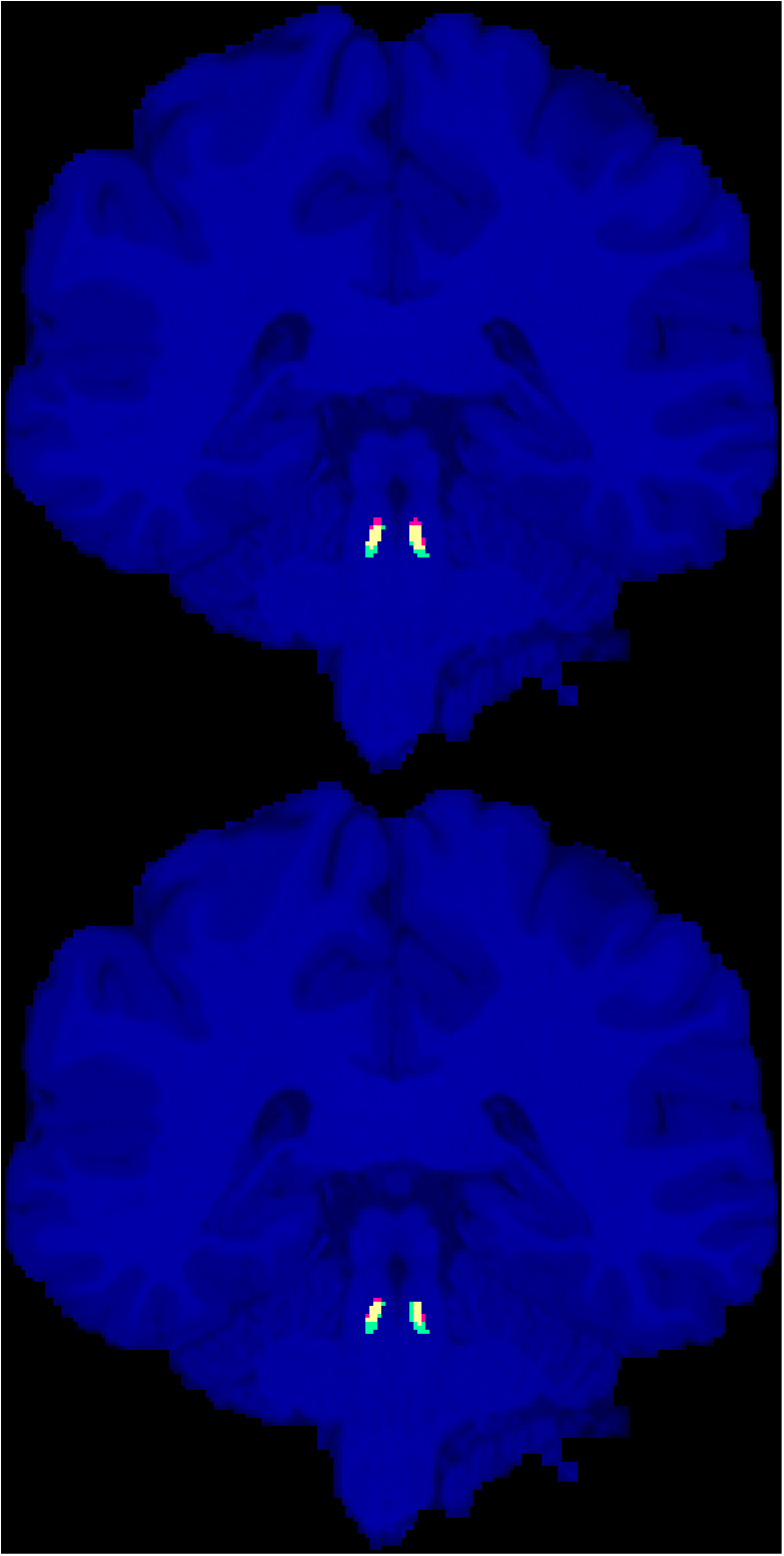
Manually annotated presumptive LC areas (green), automatic localization of LC using the ELV (top) and U-Net (bottom) methods (red), and their overlap (yellow), are shown on the coronal slice with the largest intersection with the manual LC areas for the representative HCP subject (with the median ELV Dice score).

The volume of the left and right LC regions had a cross-subject mean ± standard error of the mean (SEM) of 19.5 ± 0.8 mm^3^ and 19.8 ± 0.8 mm^3^, respectively. We also measured this for the manual labels in the external (Tona *et al* (33)) dataset; the cross-subject average volumes of the left and right LC labels were 6.9 ± 0.7 mm^3^ and 7.4 ± 0.8 mm^3^, respectively. Two-sided paired *t*-tests between the left and right volumes did not reveal a statistically significant difference between them in either dataset.

### 3.2 Automatic LC Region Estimation

We assessed our variations of the ELV and the U-Net methods (Section 2.2) via leave-one-out cross-validation on the 20-subject subset of the HCP as well as on the external dataset by Tona *et al* (33). Table 1 summarizes the median, mean, and SEM of the Dice scores.

**Table 1.** Median, mean, and standard error of the mean (SEM) of the Dice score, measuring the overlap of the automatic and manual LC regions.

| Method | Input | Cross-validation Dice score<br>(*median*)<br>(*mean* ± *SEM*) | | External validation Dice score<br>(*median*)<br>(*mean* ± *SEM*) | |
| --- | --- | --- | --- | --- | --- |
|  |  | Left LC | Right LC | Left LC | Right LC |
| ELV | phase | 0.61<br>0.61 ± 0.02 | 0.64<br>0.63 ± 0.02 | 0.02<br>0.08 ± 0.04 | 0.10<br>0.10 ± 0.03 |
| | phase + image | 0.61<br>**0.61 ± 0.02** | 0.64<br>**0.63 ± 0.02** | 0.00<br>0.00 ± 0.00 | 0.00<br>0.004 ± 0.004 |
| | phase + nmz | 0.60<br>0.61 ± 0.02 | 0.64<br>0.63 ± 0.02 | 0.00<br>0.07 ± 0.04 | 0.00<br>0.07 ± 0.03 |
| U-Net | image | 0.58<br>0.57 ± 0.03 | 0.64<br>0.60 ± 0.03 | 0.00<br>0.00 ± 0.00 | 0.00<br>0.00 ± 0.00 |
| | nmz | 0.66<br>0.61 ± 0.02 | 0.56<br>0.55 ± 0.04 | 0.14<br>0.15 ± 0.03 | 0.27<br>0.24 ± 0.03 |
| | phase | 0.63<br>0.61 ± 0.02 | 0.62<br>0.60 ± 0.02 | 0.00<br>0.03 ± 0.02 | 0.11<br>0.13 ± 0.04 |
| | image + phase | **0.62**<br>0.60 ± 0.02 | **0.64**<br>0.60 ± 0.04 | 0.00<br>0.00 ± 0.00 | 0.00<br>0.00 ± 0.00 |
| | nmz + phase | 0.62<br>0.61 ± 0.02 | 0.62<br>0.62 ± 0.02 | **0.24**<br>**0.24 ± 0.04** | **0.27**<br>**0.23 ± 0.03** |

By assessing the left and right LC together, the highest cross-validation Dice scores were achieved by ELV [phase + image] in terms of the mean (0.63 ± 0.01) and by U-Net [image + phase] in terms of the median (0.64). Figure 1 shows the labels generated by both methods for the representative subject with median ELV Dice score. For either ELV or U-Net, the best cross-validation results involved the use of the non-normalized image.

The external validation Dice scores were considerably lower, with the best mean (0.24 ± 0.03) and median (0.27) scores obtained by U-Net [nmz + phase]. Image normalization (nmz) improved the external validation Dice in all cases. The best input for either ELV or U-Net included the phase image, i.e., ELV [phase] and U-Net [nmz + phase].

Left and right LC Dice scores were significantly correlated with each other in most cross-validation results by both methods (ELV [phase + image]: r = 0.53, p = 0.02; U-Net [image]: r = 0.61, p = 0.004) and in the external validation results by ELV (ELV [phase]: r = 0.88, p = 0.0001).

### 3.3 Findings from HCP

Next, we trained a U-Net (with the default implementation, which receives the image and normalizes the patches at its first layer) on the 20 subjects and applied it to 100 HCP subjects. After correlating non-MRI variables with imaging variables (Section 2.3), only some correlations with the mean FA survived the Bonferroni correction, all of which passed the visual inspection. Table 2 lists these significant relationships, mainly with body weight and memory, and Figure 2 shows two examples. Adjusting for the intracranial volume (ICV) improved the correlation significance with the memory task accuracy, but reduced that with the body weight.

**Table 2.** Pearson’s correlation coefficient (*r*) and Bonferroni-corrected *p*-value (*p*_*B*_) of the significant correlations of the mean fractional anisotropy (FA) inside the computed LC area with non-MRI variables, without and with intracranial volume (ICV) adjustment.

| Non-MRI Variable | No ICV adjustment |  | Adjusted for ICV |  |
| --- | --- | --- | --- | --- |
|  | Left LC | Right LC | Left LC | Right LC |
| Body weight | $r = -0.52$<br>$p_B = 0.0002$ | $r = -0.52$<br>$p_B = 0.0002$ | $r = -0.44$<br>$p_B = 0.03$ | $r = -0.44$<br>$p_B = 0.02$ |
| Body mass index (BMI) | - | $r = -0.44$<br>$p_B = 0.02$ | - | $r = -0.43$<br>$p_B = 0.04$ |
| Body mass index (BMI) at heaviest time | - | $r = -0.43$<br>$p_B = 0.04$ | - | $r = -0.43$<br>$p_B = 0.046$ |
| Working memory task accuracy: 2-back place | - | $r = 0.44$<br>$p_B = 0.02$ | - | $r = 0.46$<br>$p_B = 0.009$ |

**Figure 2.**
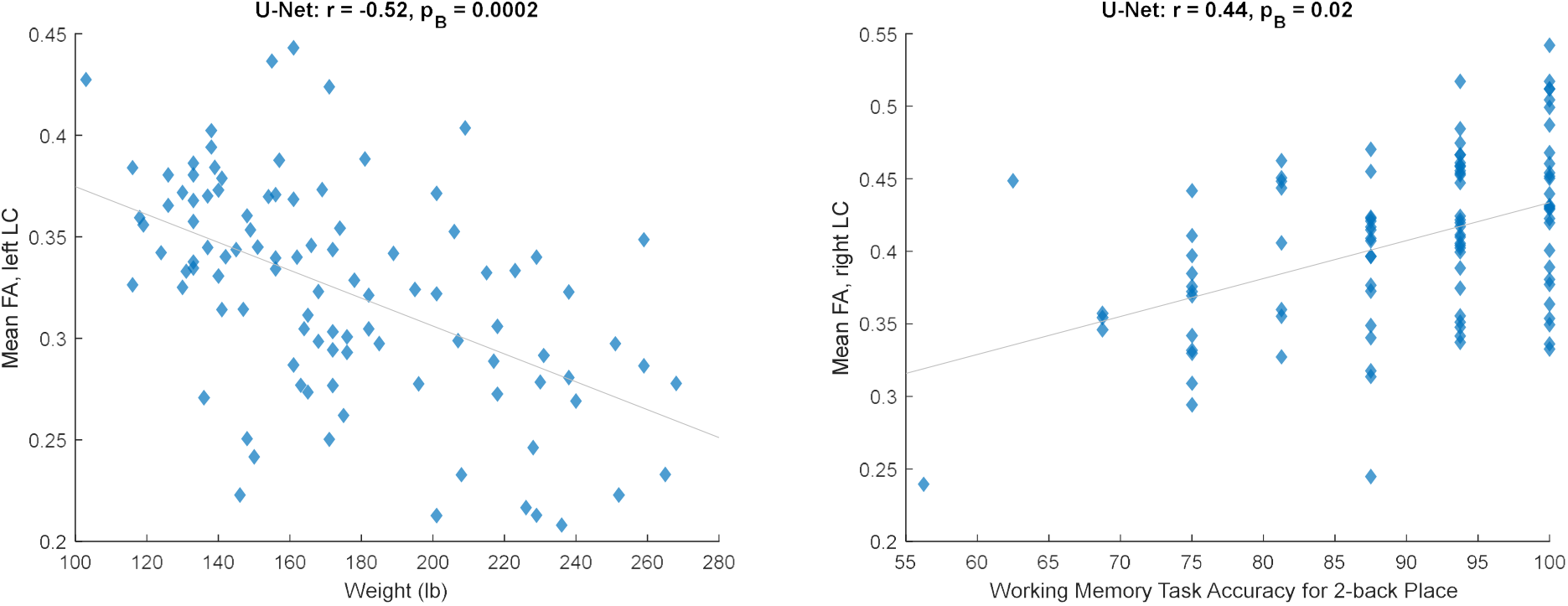
Significant relationships between the mean fractional anisotropy (FA) inside the automatically estimated LC region and non-MRI variables (corresponding to Table 2, unadjusted for ICV).

We lastly computed the strength of connectivity between each LC area and 86 other brain regions (85 + contralateral LC). After Bonferroni correction for all possible structural connections to the LC and for all non-MRI variables, none of the few significant correlations between the two passed the visual inspection (see Section 2.3).

## 4 Discussion

We have presented a new dataset of high-resolution (isotropic 0.7 mm voxel) LC areas manually approximated for 20 subjects of the HCP. This dataset is publicly available to the research community (Section 8), allowing the development of supervised tools applicable to standard 3T T_1_W MPRAGE (rather than neuromelanin-sensitive) MRI for quantitative analysis using large existing or future MRI databases. Given the lack of LC contrast on standard T_1_W images, we used dimensional information instead to estimate the region that approximately included the LC. We emphasized sensitivity for this task and created masks that were slightly larger than and contained the LC (which, if desired, could be shrunk in post-processing via the *erosion* operation). The masks had a bilateral mean volume of 19.7 ± 0.6 mm^3^, expectedly larger than the LC volume reported in the literature, such as 6.6 mm^3^ (20), 7.2 mm^3^ (that we computed from the dataset) (33), 9.5 mm^3^ (18), 12.8 mm^3^ (48), and 16.7 mm^3^ (49).

Our internal evaluation of (the optimal variations of) the ELV and U-Net automatic segmentation approaches on our data resulted in a similar cross-validation Dice score of 0.63∼0.64 for both methods. In comparison, the LC label Dice scores reported in the literature for inter-rater reliability are 0.50 (20), 0.54∼0.64 (18), 0.64 (33), and 0.68 (19), for scan-rescan reliability are 0.24∼0.48 (21) and 0.63 (50), and for automatic segmentation are 0.40 (20), 0.54∼0.64 (21), and 0.60∼0.71 (19).

Our external validation resulted in lower mean Dice scores (ELV: 0.09, U-Net: 0.24). Several reasons could account for this. Given that the average volume of our presumptive LC areas was α = 2.7 times larger than that of the LC labels in the test (Tona *et al* (33)) dataset, the Dice score between the two was capped at 0.54 (see Section 2.2.2). The cap was possibly even lower due to inter-rater variability between the two datasets, especially since our LC areas were based on dimensional information whereas the labels in the external dataset were delineated based on the LC contrast seen on neuromelanin-sensitive (3T T_1_W TSE) images. Another factor contributing to the lower Dice value may have been the domain shift, particularly caused by the different MRI field strengths of the input training (3T) and test (7T) images, which are known to reveal different MRI tissue properties (51).

We tested the hypothesis that using phase images could enhance out-of-sample segmentation. In line with our previous finding (41), the phase-only ELV map yielded a higher external-validation Dice value *without* additional modulation with the image intensity prior. The U-Net also achieved the highest accuracy in both internal and external validation when the input included the phase image. The different field strengths and acquisition protocols of the training (HCP) and test (Tona *et al* (33)) datasets may have caused inconsistencies in their image intensities. The phase image improved our external validation results by ignoring the Fourier-domain magnitude information, which possibly alleviated such inter-database inconsistencies to some extent.

The Dice scores corresponding to the approximated left and right LC regions were often significantly correlated with each other, perhaps due to their correlation with the image quality and variance of the test subject.

Following image segmentation of 100 unrelated HCP subjects and an exploratory analysis, we found significant correlations with the mean FA inside the LC region, mainly negative correlations with body weight and a positive correlation with working memory. In a similar analysis, LC connectivity was found not to be significantly correlated with non-MRI variables, possibly due to the homogeneity and narrow age range (22 – 36 years old) of the healthy HCP cohort (47). Our stringent Bonferroni correction for all compared variables and LC connections may additionally have led to type II errors (false negatives). In most related HCP studies, the LC connectivity has been measured to predefined ipsilateral target regions pertinent to disease, such as the transentorhinal cortex (15) and limbic regions (16). In contrast, we ran whole-brain tractography to explore all regions’ potential connectivity to the LC, which is especially important considering the LC’s extensive axonal branching innervating diverse remote areas throughout the brain (3).

This report has some limitations. First, the LC masks were not based on anatomical contrast and the location is approximate. Second, the LC is difficult to model given its small size. Third, partial voluming effects can introduce noticeable error in the imaging quantity derived from the LC. Should a non-MRI variable and the error in an imaging variable happen to be related to each other (e.g., by both being correlated to a third factor such as the ICV), a spurious relationship might appear in the correlation analysis. Thorough investigation of the automatically misclassified voxels to identify any potential bias is a subject of our ongoing research. For the above reasons, we do not imply our LC masks to be anatomically accurate LC labels, and caution their use where specific and accurate labels are required.

Quantitative MRI studies of the LC – requiring automatic LC region estimation – have the potential to generate imaging biomarkers for early diagnosis of neurodegenerative diseases and patient stratification (6). Making our LC neighborhood localization algorithm more specific and integrating it into FreeSurfer (42) are subjects of our future work.

## 6 Author Contributions

IA: experiment design, data analysis, writing of manuscript; JM: literature search, data curation; BF: experiment design, feedback on manuscript; JCA: manual approximation of the LC region, manuscript editing.

## 7 Funding

Support for this research was provided by the National Institutes of Health (NIH), specifically the National Institute on Aging (NIA; R56AG068261, RF1AG068261).

Additional support was provided in part by the NIA (RF1AG082223, R21AG082082, R56AG064027, R01AG064027, R01AG008122, R01AG016495, R01AG070988), the BRAIN Initiative Cell Census/Atlas Network (U01MH117023, UM1MH130981), the Brain Initiative Brain Connects consortium (U01NS132181, UM1NS132358), the National Institute for Biomedical Imaging and Bioengineering (P41EB015896, R01EB023281, R01EB006758, R21EB018907, R01EB019956, P41EB030006), the National Institute of Mental Health (R01MH121885, RF1MH123195), the National Institute for Neurological Disorders and Stroke (R01NS0525851, R21NS072652, R01NS070963, R01NS083534, U01NS086625, U24NS10059103, U24NS135561, R01NS105820), the NIH Blueprint for Neuroscience Research (U01MH093765), part of the multi-institutional Human Connectome Project, and the Michael J. Fox Foundation for Parkinson’s Research (MJFF-021226).

Computational resources were provided by the Massachusetts Life Sciences Center.

The HCP WU-Minn Consortium (U54MH091657) was funded by the 16 NIH Institutes and Centers that support the NIH Blueprint for Neuroscience Research; and by the McDonnell Center for Systems Neuroscience at Washington University.

## 8 Data Availability Statement

Upon acceptance of the paper, our manually annotated (enlarged) LC masks will be publicly available at: https://www.nitrc.org/projects/lc20

Magnetic resonance images were provided by:

- the Human Connectome Project (HCP, RRID:SCR_006942) (39), WU-Minn Consortium (Principal Investigators: David Van Essen and Kamil Ugurbil): https://www.humanconnectome.org/study/hcp-young-adult
- Klodiana-Daphne Tona *et al* (33): https://doi.org/10.34894/PMQHZD

The following toolboxes were used for data processing and analysis:

- Our MATLAB (RRID:SCR_001622) toolboxes for:
  - the expected label value (ELV) supervised image segmentation (41): https://www.nitrc.org/projects/elv
  - the reconstruction of the orientation distribution function in constant solid angle (44), Hough-transform tractography (45), and connectivity matrix computation and augmentation (46): http://www.nitrc.org/projects/csaodf-hough
- FreeSurfer (RRID:SCR_001847) (42): https://freesurfer.net
- SPM12 (RRID:SCR_007037) (43): https://www.fil.ion.ucl.ac.uk/spm/software/spm12

